# Strongly bactericidal all-oral β-lactam combinations for the treatment of *Mycobacterium abscessus* lung disease

**DOI:** 10.1101/2022.06.07.495238

**Authors:** Dereje A. Negatu, Matthew D. Zimmerman, Véronique Dartois, Thomas Dick

## Abstract

Bioactive forms of oral β-lactams were screened in vitro against *Mycobacterium abscessus* with and without the bioactive form of the oral β-lactamase inhibitor avibactam-ARX1796. Sulopenem was equally active without avibactam, while tebipenem, cefuroxime and amoxicillin required avibactam for optimal activity. Systematic pairwise combination of the four β-lactams revealed strong bactericidal synergy for each of sulopenem, tebipenem and cefuroxime combined with amoxicillin in the presence of avibactam. These all-oral β-lactam combinations warrant clinical evaluation.

## MAIN TEXT

*M. abscessus* lung disease is treated with an oral macrolide (clarithromycin (CLR) or azithromycin) in combination with several, largely underperforming antibiotics including parenteral amikacin, one of the two parenteral β-lactams imipenem (IPM) or cefoxitin (FOX), and tigecycline (1). Patients are often treated for years, until sputum cultures remain negative for 12 months, if culture conversion is achieved. Chemotherapies, complicated by the need to use injectable drugs, are not only long but often toxic (1, 2). Treatment is further tangled by widespread inducible resistance to the oral macrolide component due to the presence of the ribosome methylase gene *erm41* (3–5). In short, there is no reliable cure for *M. abscessus* lung disease. Novel, well tolerated, bactericidal, and importantly, oral treatment options are sorely needed (6, 7).

β-lactams are bactericidal and display overall excellent tolerability profiles (8). However, IPM and FOX, the standard of care carbapenem and cephalosporin, are administered intravenously, limiting their clinical utility given the very long treatment duration required to control or cure *M. abscessus* lung disease. They also show modest in vitro activity (9, 10), leading to poor pharmacokinetic-pharmacodynamic target attainment compared to those achieved against other bacterial infections.

Different classes of β-lactams, and different members within a class, differentially inhibit the numerous *M. abscessus* transpeptidases and other enzymes involved in peptidoglycan synthesis (11–16). Thus, multiple recent reports have demonstrated the potential of combining two β-lactams to achieve additive and synergistic effects in vitro (17–21) as well as in vivo (22).

A number of oral β-lactams are in clinical development or in clinical use for other bacterial infections (23, 24). Furthermore, an oral form (ARX1796) of the β-lactamase inhibitor avibactam (AVI), inhibiting the major β-lactamase MAB_2875 of *M. abscessus* (9, 25), recently entered clinical development (26) (ClinicalTrials.gov Identifier: NCT03931876).

Here, our goal was to identify oral β-lactam pairs that exert synergistic bactericidal activity (with or without oral AVI) and can be repurposed to treat *M. abscessus* lung disease. First, a collection of the bioactive forms of 22 oral β-lactams, including penems, carbapenems, cephalosporins and penicillins, was screened at a single concentration of 12.5 μM with and without the bioactive form of AVI at a fixed concentration of 4 μg/ml (14 μM) to identify antibiotics with attractive anti-*M. abscessus* growth inhibitory activity (27). Growth inhibition against the type strain *M. abscessus* susbsp. *abscessus* ATCC19977 was measured in 7H9 broth using OD_600_ as readout (28). Applying > 80 % growth inhibition as cut-off, we identified one penem (sulopenem, SUP), one carbapenem (tebipenem, TBP), one cephalosporin, (cefuroxime, CXM) and two penicillins (ampicillin (AMP) and amoxicillin (AMX)) as active agents. At 12.5 μM, SUP and CXM were equally active with and without AVI. TBP, AMP and AMX required AVI for activity (Figure 1, Figure S1).

**Fig. 1.**
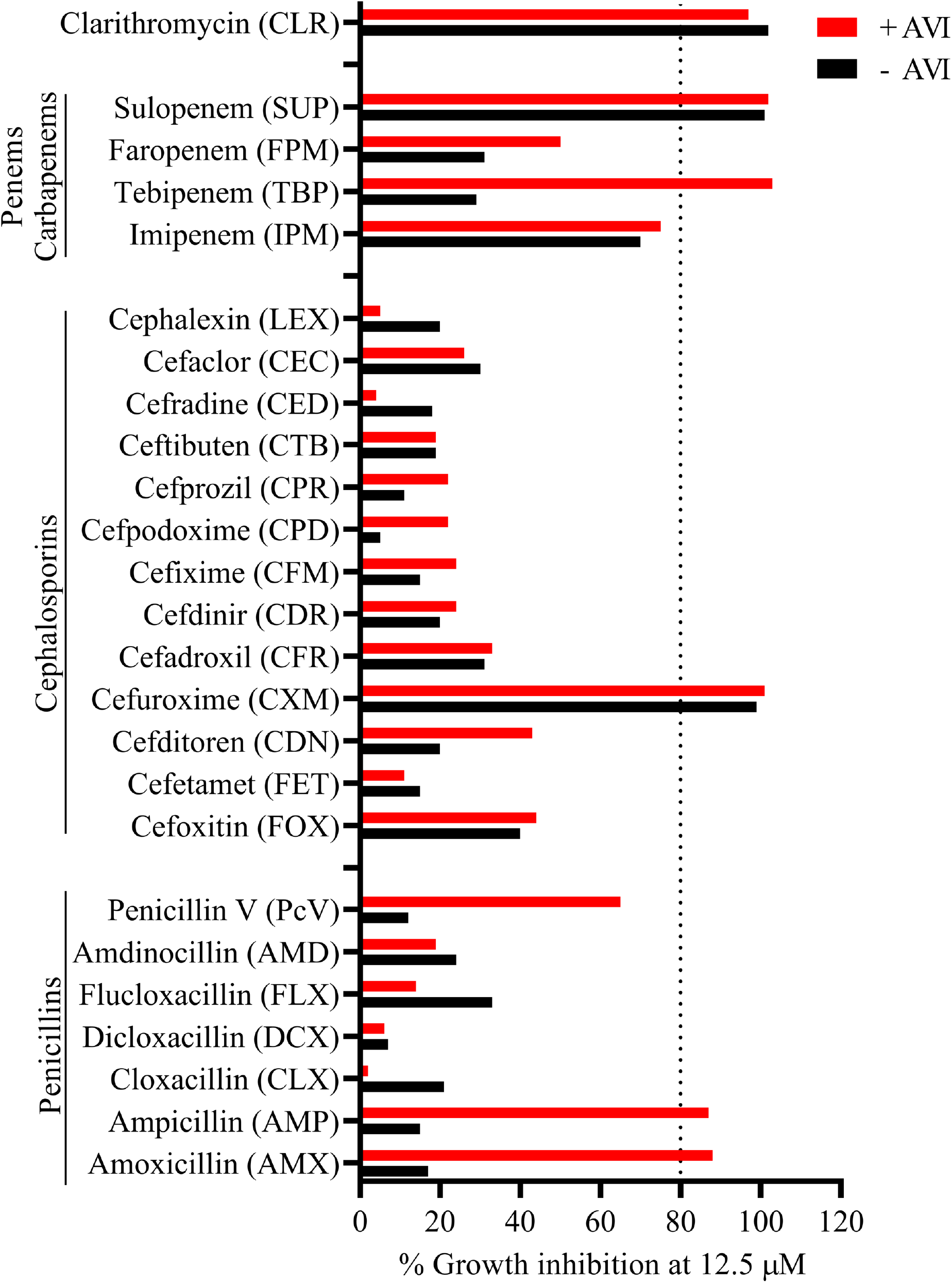
Single point growth inhibition screen of β-lactams with and without 4 μg/mL AVI against *M. abscessus* ATCC19977. A collection of the bioactive forms of 22 oral β-lactams was screened at 12.5 μM. Percent growth inhibition is shown. Dashed line, 80% growth inhibition. CLR was included as positive control, IPM and FOX as clinically used parenteral comparators. The experiment was carried out twice yielding similar results. Compound sources, oral prodrug forms (if applicable) and clinical status are described in Table S1.

To confirm the results from the single point screen, we determined the minimum inhibitory concentration (MIC) of SUP, TBP, CXM, and AMX (the early generation penicillin AMP was not followed up) with or without AVI (4 μg/mL). MIC was defined as 90% growth inhibition and derived from dose response curves (28) (Table 1, Figure S2). The MIC of SUP was 2.5 μM and AVI-independent. TBP had an MIC of 4 μM in the presence of AVI. CXM showed a weak AVI dependency, with MICs of 5 μM and 10 μM with and without AVI, respectively. AMX exhibited a unique activity profile with a modest MIC of 25 μM in the presence of AVI but a substantially lower MIC_50_ (concentration inhibiting 50% of growth) of 3 μM, similar to the MIC_50_ of SUP, TBP and CXM (Figure S2). The two injectable comparators IPM and FOX (which are both AVI independent (9)) showed MICs of 20 and 30 μM as reported (9). (Table 1, Figure S2).

**Table 1.** Activity of SUP, TBP, CXM and AMX without and with 4 μg/mL AVI against *M. abscessus* complex strains.^*a*^

| *M. abscessus* strain | *erm41*<sup>d</sup><br>sequevar | CLR<br>susceptibility | SUP |  | TBP |  | CXM |  | AMX |  | IPM<sup>b</sup> | FOX<sup>b</sup> | CLR<sup>c</sup> |
| --- | --- | --- | --- | --- | --- | --- | --- | --- | --- | --- | --- | --- | --- |
|  |  |  | MIC | MIC+ | MIC | MIC+ | MIC | MIC+ | MIC | MIC+ |  |  |  |
| Reference strains |  |  |  |  |  |  |  |  |  |  |  |  |  |
| subsp. *abscessus* ATCC19977 | T28 | Resistant | 2.5 | 2.0 | 25.0 | 4.0 | 10.0 | 5.0 | >100 | 25.0<sup>a</sup> | 20.0 | 30.0 | 1.6 |
| subsp. *bolletii* CCUG50184T | T28 | Resistant | 3.5 | 2.0 | 30.0 | 5.0 | 20.0 | 10.0 | >100 | 40.0 | 30.0 | 30.0 | 5.0 |
| subsp. *massiliense* CCUG48898T | Deletion | Sensitive | 7.0 | 5.0 | 40.0 | 7.0 | 30.0 | 10.0 | >100 | 100.0 | 40.0 | 45.0 | 0.4 |
| Clinical isolates<sup>e</sup> |  |  |  |  |  |  |  |  |  |  |  |  |  |
| subsp. *abscessus* Bamboo | C28 | Sensitive | 3.0 | 2.5 | 25.0 | 4.0 | 10.0 | 8.0 | >100 | 40.0 | 20.0 | 35.0 | 0.4 |
| subsp. *abscessus* K21 | C28 | Sensitive | 6.3 | 5.0 | 40.0 | 4.0 | 30.0 | 20.0 | >100 | 100.0 | 25.0 | 40.0 | 0.5 |
| subsp. *abscessus* M9 | T28 | Resistant | 3.0 | 2.5 | 35.0 | 3.5 | 10.0 | 5.0 | >100 | 40.0 | 15.0 | 35.0 | 2.5 |
| subsp. *abscessus* M199 | T28 | Resistant | 3.0 | 2.5 | 25.0 | 3.0 | 12.5 | 8.0 | >100 | 75.0 | 20.0 | 35.0 | 6.0 |
| subsp. *abscessus* M337 | T28 | Resistant | 2.0 | 2.2 | 30.0 | 3.0 | 10.0 | 8.0 | >100 | 60.0 | 15.0 | 30.0 | 3.0 |
| subsp. *abscessus* M404 | C28 | Sensitive | 3.5 | 2.5 | 30.0 | 3.5 | 20.0 | 7.0 | >100 | 40.0 | 20.0 | 35.0 | 0.4 |
| subsp. *abscessus* M422 | T28 | Resistant | 2.5 | 2.0 | 25.0 | 3.0 | 10.0 | 5.0 | >100 | 40.0 | 12.5 | 35.0 | 1.5 |
| subsp. *bolletii* M232 | T28 | Resistant | 2.5 | 2.0 | 40.0 | 3.5 | 15.0 | 8.0 | >100 | 50.0 | 15.0 | 40.0 | 2.0 |
| subsp. *bolletii* M506 | C28 | Sensitive | 2.5 | 2.0 | 30.0 | 3.5 | 15.0 | 8.0 | >100 | 70.0 | 18.0 | 35.0 | 0.4 |
| subsp. *massiliense* M111 | Deletion | Sensitive | 4.0 | 3.5 | 35.0 | 4.0 | 15.0 | 10.0 | >100 | 100.0 | 30.0 | 35.0 | 0.4 |

<sup>a</sup> Cultures were treated with increasing concentrations of β lactams without ('MIC') or with 4 µg/ml AVI ('MIC+') (27). Values present the concentrations that achieved 90% inhibition of growth and are the means of three independent experiments. Note that AMX + AVI achieved 80% inhibition of growth at ~10 µM (Figure S2).
<sup>b</sup> The clinically used parenteral comparators IPM and FOX were only tested alone as AVI does not affect activity of these β lactams (Figure 1) (9).
<sup>c</sup> CLR, assay control. Note increased MIC values for CLR resistant strains.
<sup>d</sup> *erm41*, ribosome methylase gene conferring inducible CLR resistance. ‘C28’ and ‘deletion’ sequevars are inactive *erm41* alleles and susceptible to CLR. The ‘T28’ sequevar is functional and confers inducible resistance to CLR (3). <sup>e</sup> *M. abscessus* Bamboo (37), K21 (38) and M strains (39) were reported previously.

To confirm the growth inhibitory activity of SUP, TBP+AVI, CXM+AVI and AMX+AVI in an orthogonal assay, we determined their MIC against *M. abscessus* ATTC19977 using the agar dilution method (29) and found agar MICs in the range of broth MICs (Figure S3), with the exception of AMX which interestingly had a lower agar than liquid MIC (6 μM versus 25 μM).

To determine whether the attractive activity of SUP, TBP+AVI, CXM+AVI and AMX+AVI against the type strain *M. abscessus* susbsp. *abscessus* ATCC19977 was retained against the broader *M. abscessus* complex (30), broth MICs were measured against the reference strains of the two other subspecies *M. abscessus* subsp. *bolletii* CCUG50184T and *M. abscessus* subsp. *massiliense* CCUG48898T, as well as a panel of clinical isolates, including *erm41* macrolide resistant strains (Table 1, Figure S2). Potency of the active β-lactams was largely comparable across the three subspecies of the *M. abscessus* complex. Again, CXM activity was 2 to 3-fold enhanced in the presence of AVI and AMX+AVI displayed modest MIC_90_ (Table 1), but substantially lower (3 – 4 μM) MIC_50_ (Figure S2).

Next, we measured the bactericidal activity in dose response time-kill experiments (28). *M. abscessus* ATCC19977 cultures were grown in 7H9 and treated with MIC multiples of the β-lactams for 5 days (in the presence of 4 μg/mL AVI when required), and CFU were measured by plating samples on 7H10 agar. All four β-lactams achieved pronounced reductions in viable counts, up to 4-log CFU reduction after 3 days of incubation with 8-fold MIC (Figure 2A). Interestingly, regrowth was observed in most cultures between day 3 and 5 for the oral β-lactams, and even earlier for the injectables, which we hypothesized was associated with the limited aqueous stability of β-lactams (31, 32). To test the potential decay hypothesis, drug concentrations were followed in 7H9 medium over 5 days using high-pressure liquid chromatography coupled to tandem mass spectrometry (LC-MS/MS) (33). The lactam ring-containing drugs (i.e., all study drugs but not AVI) were unstable to various extents, mostly in line with the extent and timing of regrowth of the cultures (Figure 2B). Half-lives (t ½) of the β-lactams in 7H9 ranged from ∼0.5 days for the markedly unstable IMP used as comparator (32) to ∼5 days for the most stable SUP (Figure 2B). Since β-lactams also undergo spontaneous and enzymatic hydrolysis in plasma (34), we measured stability of the four oral β-lactams and AVI in mouse plasma to determine which pair would be most suitable for in vivo efficacy in murine models of *M. abscessus* infection. We found that TBP, AMX and AVI have a plasma t ½ ≥ 24h (Figure 2C), indicating that the TBP/AMX+AVI combinations could be prioritized as a case study to determine how these in vitro bactericidal synergies translate in vivo.

**Fig. 2.**
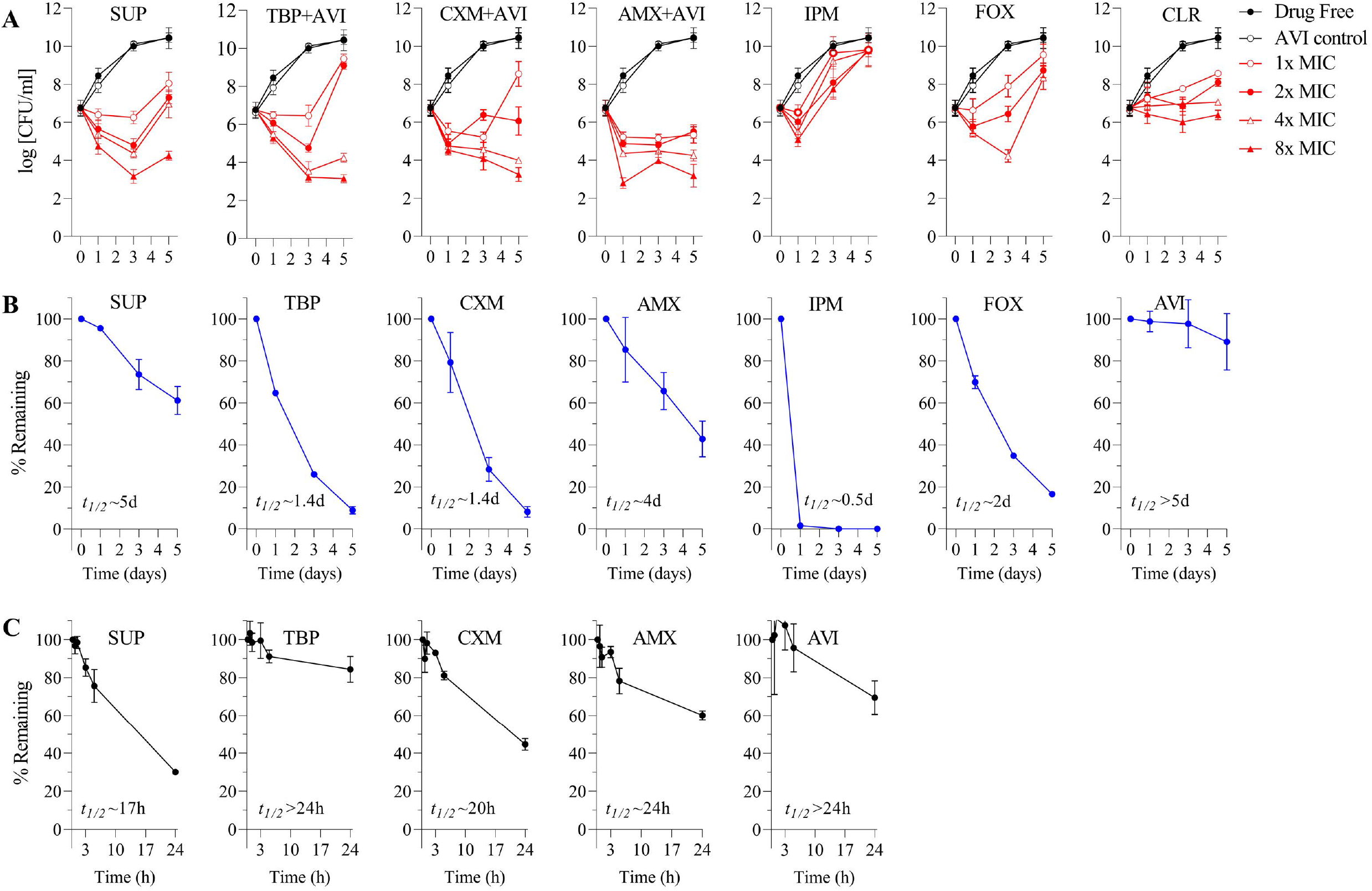
Dose response time-kill curves of SUP, TBP+AVI, CXM+AVI and AMX-AVI against *M. abscessus* ATCC19977 and drug stability in culture medium and mouse plasma. (A) Time concentration kill curves. Cultures of *M. abscessus* ATCC19977 were treated with MIC (Table 1) and multiples of MIC of SUP (alone), TBP, CXM, AMX (in combination with 4 ug/mL AVI) for 5 days and viability of the cultures was monitored by CFU determination. CLR was included as assay control. IPM and FOX were included as clinically used parenteral comparators. (B) Stability of the β-lactams tested in (A) and AVI in 7H9 broth over a 5-day incubation period at 37ºC. Percent remaining was calculated relative to time 0 concentration (10 μM). Half-life was estimated from the decay curves. (C) Mouse plasma stability of the four oral β-lactams and AVI over 1-day incubation period. Experiments in (A) were performed twice independently, generating similar data, and one representative set is shown. Experiments in (B) and (C) were carried out twice independently and means and standard deviations are shown.

Taken together, these results confirm and extend prior studies showing attractive growth inhibitory and bactericidal anti-*M. abscessus* activity of SUP (35), TBP+AVI (27), CXM+AVI (9) and AMX+AVI (9), suggesting them as repurposing candidates.

To determine potential growth inhibition synergies of the four β-lactams, systematic pairwise checkerboard analyses were carried out with *M. abscessus* ATCC19977 (36). AVI was included at 4 μg/mL in all assays as at least one partner of each dual combination requires the β-lactamase inhibitor (Table 2). Interestingly, the three AMX-containing pairs were synergistic, while the other three were additive (Table 2). Synergistic activity of the AMX-containing pairs was confirmed in checkerboard assays against the broader *M. abscessus* complex and the clinical isolate collection. TBP and CXM combined with AMX retained strong synergistic activity against all tested strains and isolates, while SUP+AMX was additive against some of the isolates (Table 3).

**Table 2.** Checkerboard growth inhibition analysis of pairwise combinations of SUP, TBP, CXM, and AMX in the presence of 4 μg/mL AVI against *M. abscessus* ATCC19977.

| β-lactams | MIC [µM] |  | FICI<sup>c</sup> | Interpretation<sup>d</sup> |
| --- | --- | --- | --- | --- |
|  | Alone<sup>a</sup> | Comb<sup>b</sup> |  |  |
| SUP<br>TBP | 4.0<br>2.5 | 0.5<br>1.2 | 0.61 | Additive |
| SUP<br>CXM | 4.0<br>5.0 | 0.8<br>2.0 | 0.60 | Additive |
| SUP<br>AMX | 4.0<br>25.0 | 0.5<br>6.0 | 0.37 | Synergy |
| TBP<br>CXM | 2.5<br>5.0 | 1.5<br>1.0 | 0.80 | Additive |
| TBP<br>AMX | 2.5<br>25.0 | 0.5<br>6.0 | 0.44 | Synergy |
| CXM<br>AMX | 5.0<br>25.0 | 0.8<br>3.0 | 0.28 | Synergy |

<sup>a</sup> MIC of single drugs, with 4 µg/mL AVI in the case of TBP, CXM and AMX.
<sup>b</sup> MIC of the combination (all in the presence of 4 µg/mL AVI as at least one partner drugs require AVI for activity).
<sup>c</sup> Fractional Inhibitory Concentration Index, calculated using the concentration at which at least 90% growth inhibition of the cultures was observed. $FICI = [(concentration\ of\ drug\ A\ in\ combination / concentration\ of\ drug\ A\ alone) + (concentration\ of\ drug\ B\ in\ combination / concentration\ of\ drug\ B\ alone)]$ .
<sup>d</sup> FICI, ≤ 0.5, synergistic; 0.5 to 1.0, additive; > 1.0 to < 2, indifferent; ≥ 2.0, antagonistic (40).
The experiment was repeated once yielding similar results.

**Table 3.** Checkerboard growth inhibition analysis of SUP+AMX, TBP+AMX and CXM+AMX in the presence of 4 μg/mL AVI against *M. abscessus* complex strains.

| *M. abscessus* strain | SUP+AMX |  |  |  |  | TBP+AMX |  |  |  |  | CXM+AMX |  |  |  |  |
| --- | --- | --- | --- | --- | --- | --- | --- | --- | --- | --- | --- | --- | --- | --- | --- |
|  | MIC [µM]<sup>a</sup> |  |  |  | FICI<sup>b</sup> | MIC [µM]<sup>a</sup> |  |  |  | FICI<sup>b</sup> | MIC [µM]<sup>a</sup> |  |  |  | FICI<sup>b</sup> |
|  | SUP Alone | AMX Alone | SUP Comb | AMX Comb |  | TBP Alone | AMX Alone | TBP Comb | AMX Comb |  | CXM Alone | AMX Alone | CXM Comb | AMX Comb |  |
| **Reference strains** |  |  |  |  |  |  |  |  |  |  |  |  |  |  |  |
| subsp. *abscessus* ATCC19977 | 2.5 | 25.0 | 0.5 | 7.0 | 0.48 | 4.0 | 25.0 | 0.4 | 7.0 | 0.38 | 6.3 | 25 | 0.5 | 4.0 | 0.24 |
| subsp. *bolletii* CCUG50184T | 2.5 | 50 | 0.7 | 10.0 | 0.48 | 4.0 | 50.0 | 1.0 | 4.0 | 0.33 | 8.0 | 50.0 | 1.5 | 6.3 | 0.31 |
| subsp. *massiliense* CCUG48898T | 4.0 | 100 | 1.0 | 25.0 | 0.50 | 8.0 | 100.0 | 1.0 | 10.0 | 0.22 | 12.5 | 100.0 | 1.5 | 25 | 0.37 |
| **Clinical isolates** |  |  |  |  |  |  |  |  |  |  |  |  |  |  |  |
| subsp. *abscessus* Bamboo | 2.5 | 50.0 | 0.5 | 10.0 | 0.40 | 4.5 | 40.0 | 0.5 | 5.0 | 0.24 | 8.0 | 50.0 | 1.5 | 5.0 | 0.29 |
| subsp. *abscessus* K21 | 5.0 | 100.0 | 1.0 | 25.0 | 0.45 | 5.0 | 100.0 | 1.5 | 12.5 | 0.43 | 12.5 | 100.0 | 1.5 | 6.3 | 0.18 |
| subsp. *abscessus* M9 | 2.5 | 50.0 | 0.5 | 10.0 | 0.40 | 4.0 | 50.0 | 0.8 | 6.3 | 0.33 | 6.3 | 50.0 | 0.8 | 12.5 | 0.38 |
| subsp. *abscessus* M199 | 2.5 | 100.0 | 0.5 | 25.0 | 0.45 | 4.5 | 80.0 | 0.5 | 10.0 | 0.24 | 8.0 | 100.0 | 1.5 | 10.0 | 0.29 |
| subsp. *abscessus* M337 | 2.0 | 75.0 | 0.4 | 25.0 | 0.53 | 4.5 | 75.0 | 0.8 | 12.5 | 0.34 | 8.0 | 75.0 | 1.5 | 10.0 | 0.32 |
| subsp. *abscessus* M404 | 2.0 | 50.0 | 0.4 | 12.5 | 0.45 | 4.5 | 50.0 | 0.8 | 6.3 | 0.30 | 8.0 | 50.0 | 1.0 | 5.0 | 0.23 |
| subsp. *abscessus* M422 | 2.5 | 50.0 | 0.5 | 10.0 | 0.40 | 4.5 | 50.0 | 0.5 | 6.0 | 0.23 | 8.0 | 50.0 | 1.5 | 5.0 | 0.29 |
| subsp. *bolletii* M232 | 2.5 | 50.0 | 0.8 | 12.5 | 0.57 | 4.5 | 50.0 | 1.0 | 6.3 | 0.35 | 8.0 | 50.0 | 2.0 | 6.3 | 0.38 |
| subsp. *bolletii* M506 | 2.5 | 75.0 | 0.5 | 15.0 | 0.40 | 4.5 | 75.0 | 0.5 | 10.0 | 0.24 | 8.0 | 80.0 | 1.5 | 10.0 | 0.31 |
| subsp. *massiliense* M111 | 2.5 | 100.0 | 0.8 | 25.0 | 0.57 | 4.5 | 75.0 | 0.8 | 12.5 | 0.34 | 8.0 | 100.0 | 1.5 | 8.0 | 0.27 |

<sup>a</sup> Alone and Comb as described in Table 2.
<sup>b</sup> FICI, ≤ 0.5, synergistic; 0.5 to 1.0, additive (40).

To determine whether the three synergistic β-lactam pairs also exerted potentiation of bactericidal activity, time-kill experiments were carried out with *M. abscessus* ATCC19977 in 7H9 and the effect of treatment on viability was measured by plating on 7H9 agar (28). To uncover potential bactericidal synergy, we combined SUP, TBP or CXM at their MIC, concentrations that achieve little bactericidal effect (Figure 2A), with AMX at 10 μM (at which the drug inhibits 80 % growth, Figure S2)) and 4 μg/mL AVI. Impressively, each of the three combinations achieved more than 4-log reduction in viable counts after 3 days of treatment (Figure 3). In comparison, 8x MIC of each individual β-lactam was required to achieve a similar degree of killing (Figure 2). Further reducing AMX concentration to 5 or 2.5 μM still achieved a 4-log reduction after 5 days of treatment, reenforcing the notion that AMX strongly potentiates the bactericidal activity of SUP, TBP and CXM. In addition, the combinations not only killed effectively at lower concentrations than individual β lactams, they also prevented the re-growth observed in cultures treated with single drugs (Figure 2A).

**Fig. 3.**
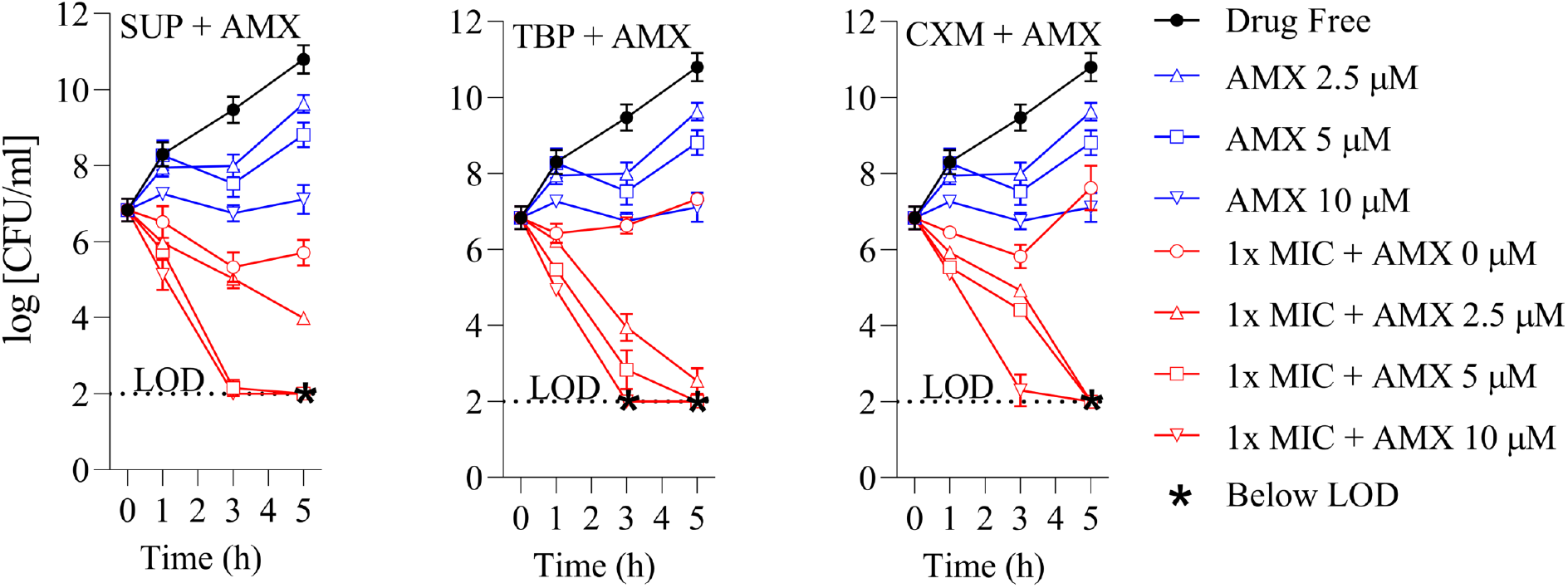
Time**-**kill curves of SUP+AMX, TBP+AMX and CXM+AMX in the presence of 4 μg/mL AVI against *M. abscessus* ATCC19977. Red curves: Cultures of *M. abscessus* ATCC19977 were treated with 1x MIC of SUP, TBP or CXM (Table 1) with or without 10, 5 or 2.5 μM AMX in the presence of 4 μg/mL AVI for 5 days. Note, 10 μM AMX suppresses growth by 80% (Figure S2). Bacterial viability was monitored by CFU determination. Blue curves: treatment of cultures with 10, 5, 2.5 μM AMX alone in the presence of 4 μg/mL AVI. LOD, limit of detection (100 CFU/mL). The experiment was carried out twice independently, generating similar results, and one representative set of plots is shown.

In conclusion, four oral β-lactams SUP, TBP, CXM and AMX were identified as bactericidal against *M. abscessus* at clinically achievable concentrations. TBP, CXM and AMX required the β-lactamase inhibitor AVI for optimal activity whereas SUP’s activity was AVI independent. Pairwise combinations revealed three novel triple combinations (SUP or TBP or CXM with AMX plus AVI) showing both bacteriostatic and bactericidal synergy. Interestingly, all three β-lactam pairs contained AMX, which preferentially targets *M. abscessus* D,D-carboxypeptidase (14), whereas the carbapenem TBP was shown to inhibit L,D-transpeptidases Ldt_Mab1_ and Ldt_Mab2_ (13) and D,D-transpeptidases PonA1, PonA2, and PbpA (11). The specific targets of SUP and CXM have not been identified. However, similar to TBP, other penems and cephalosporins were also shown to preferentially target L,D- and D,D-transpeptidases (11–13). The differential inhibition of D,D-carboxypeptidase by AMX and of L,D- and D,D-transpeptidases by TBP and possibly SUP and CXM may provide the mechanistic basis for the observed synergistic effects of the AMX-containing β-lactam couples, since they would inhibit different enzymes of the same cellular process, i.e. peptidoglycan synthesis. Oral forms of SUP and TBP as well as AVI are currently in clinical development for other diseases and oral CXM and AMX are approved drugs (Table S1). Thus, the compounds and combinations identified in this study present drug candidates that can enter clinical development for *M. abscessus* lung disease.

## ACKNOWLEDGMENTS

We are grateful to Wei Chang Huang (Taichung Veterans General Hospital, Taichung, Taiwan) for providing *M. abscessus* Bamboo, to Jeanette W.P. Teo (Department of Laboratory Medicine, National University Hospital, Singapore) for providing the collection of *M. abscessus* clinical M isolates, and to Sung Jae Shin (Department of Microbiology, Yonsei University College of Medicine, Seoul, South Korea) and Won-Jung Koh (Division of Pulmonary and Critical Care Medicine, Samsung Medical Center, Seoul, South Korea) for providing *M. abscessus* K21. We thank Rubén González del Río and Mónica Cacho-Izquierdo (Global Health Pharma Unit, GlaxoSmithKline, Tres Cantos, Madrid, Spain) for valuable discussion. Research reported in this work was supported by the National Institute of Allergy and Infectious Diseases of the National Institutes of Health under Award Number R01AI132374. The content is solely the responsibility of the authors and does not necessarily represent the official views of the National Institutes of Health.

## AUTHOR CONTRIBUTIONS

Investigation: D.A.N., M.D.Z.; Writing – Original Draft: D.A.N., T.D.; Writing – Review & Editing: all authors; Funding Acquisition: T.D.; Supervision: V.D., T.D.

## CONFLICT OF INTEREST STATEMENT

The authors declare no commercial or financial relationships that could be construed as a potential conflict of interest.

## Notes

### Competing Interest Statement

The authors have declared no competing interest.

### Summary of Updates

supplemental materials added

