## Supplemental Table and Figures for "Strongly bactericidal all-oral β-lactam combinations for the treatment of *Mycobacterium abscessus* lung disease"

Running Title: Combination of oral β lactams against *M. abscessus*

Keywords: Non-tuberculous mycobacteria, NTM, synergy, sulopenem, tebipenem, cefuroxime, amoxicillin, avibactam

### SUPPLEMENTAL TABLES

**Table S1.** Drugs used in the study: oral prodrug form, source, solvent and clinical status

| No. | Drug | Prodrug form | Class | Catalog # | Source | Solvent | FDA status | Clinical development |
|---|---|---|---|---|---|---|---|---|
| 1 | Clarithromycin (CLR) | N/A | Macrolide | C9742 | Sigma-Aldrich | DMSO | Approved | |
| 2 | Avibactam (AVI) | ARX-1796 | Diazabicyclooctane | HY-14879A | MedChemExpress | DMSO | Approved[a] | [a] Phase 1 (NCT03931876) |
| 3 | Sulopenem (SUP) | Sulopenem etzadroxil | Penem | PZ0042 | Sigma-Aldrich | DMSO | Not approved | [b] Phase 3 (NCT03357614) |
| 4 | Faropenem (FPM) | Faropenem medoxomil | Penem | F8182 | Sigma-Aldrich | DMSO | Not approved | [c] Phase 2 (NCT02381470) |
| 5 | Tebipenem (TBP) | Tebipenem pivoxil | Carbapenem | 161715-21-5 | MuseChem | DMSO | Not approved | [d] Phase 3 (NCT03788967) |
| 6 | Imipenem (IPM) | N/A | Carbapenem | PHR1796 | Sigma-Aldrich | Water | Approved | |
| 7 | Cephalexin (LEX) | N/A | Cephalosporin | PHR1848 | Sigma-Aldrich | Water | Approved | |
| 8 | Cefaclor (CEC) | N/A | Cephalosporin | PHR1283 | Sigma-Aldrich | Water | Approved | |
| 9 | Cefradine (CED) | N/A | Cephalosporin | C0690000 | Sigma-Aldrich | DMSO | Approved | |
| 10 | Ceftibuten (CTB) | N/A | Cephalosporin | SML0037 | Sigma-Aldrich | DMSO | Approved | |
| 11 | Cefprozil (CPR) | N/A | Cephalosporin | Y0001371 | Sigma-Aldrich | DMSO | Approved | |
| 12 | Cefpodoxime (CPD) | Cefpodoxime proxetil | Cephalosporin | 32344 | Sigma-Aldrich | DMSO | Approved | |
| 13 | Cefixime (CFM) | N/A | Cephalosporin | CDS021590 | Sigma-Aldrich | DMSO | Approved | |
| 14 | Cefdinir (CDR) | N/A | Cephalosporin | C7118 | Sigma-Aldrich | DMSO | Approved | |
| 15 | Cefadroxil (CFR) | N/A | Cephalosporin | C0650000 | Sigma-Aldrich | DMSO | Approved | |
| 16 | Cefuroxime (CXM) | Cefuroxime axetil | Cephalosporin | C4417 | Sigma-Aldrich | DMSO | Approved | |
| 17 | Cefditoren (CDN) | Cefditoren pivoxil | Cephalosporin | HY-17452 | MedChemExpress | DMSO | Approved | |
| 18 | Cefetamet (FET) | Cefetamet pivoxil | Cephalosporin | HY-B1894A | MedChemExpress | DMSO | Not approved | [e] Phase 4 (NCT04664803) |
| 19 | Cefoxitin (FOX) | N/A | Cephalosporin | C4786 | Sigma-Aldrich | DMSO | Approved | |
| 20 | Penicillin V (PcV) | N/A | Penicillin | PHR2644 | Sigma-Aldrich | DMSO | Approved | |
| 21 | Amdinocillin (AMD) | Pivmecillinam | Penicillin | 32887-01-7 | MuseChem | DMSO | Approved | |
| 22 | Flucloxacillin (FLX) | N/A | Penicillin | SML1023 | Sigma-Aldrich | DMSO | Approved | |
| 23 | Dicloxacillin (DCX) | N/A | Penicillin | 46182 | Sigma-Aldrich | DMSO | Approved | |
| 24 | Cloxacillin (CLX) | N/A | Penicillin | PHR1922 | Sigma-Aldrich | DMSO | Approved | |
| 25 | Ampicillin (AMP) | Bacampicillin | Penicillin | HY-B0522 | MedChemExpress | Water | Approved | |
| 26 | Amoxicillin (AMX) | N/A | Penicillin | 1031503 | Sigma-Aldrich | DMSO | Approved | [f] Phase 2 (NCT02381470) |

[a] Only the injectable form of AVI is approved. The oral AVI prodrug ARX-1796 is in Phase 1 clinical development. [b] Sulopenem etzadroxil, in clinical development for complicated urinary tract infections. [c] Faropenem, in clinical development for tuberculosis. [d] Tebipenem pivoxil hydrobromide, in clinical development for complicated urinary tract infections and acute pyelonephritis. [e] Cefetamet pivoxil, in clinical development for sinusitis. [f] Amoxicillin, in clinical development for tuberculosis.

### SUPPLEMENTAL FIGURES

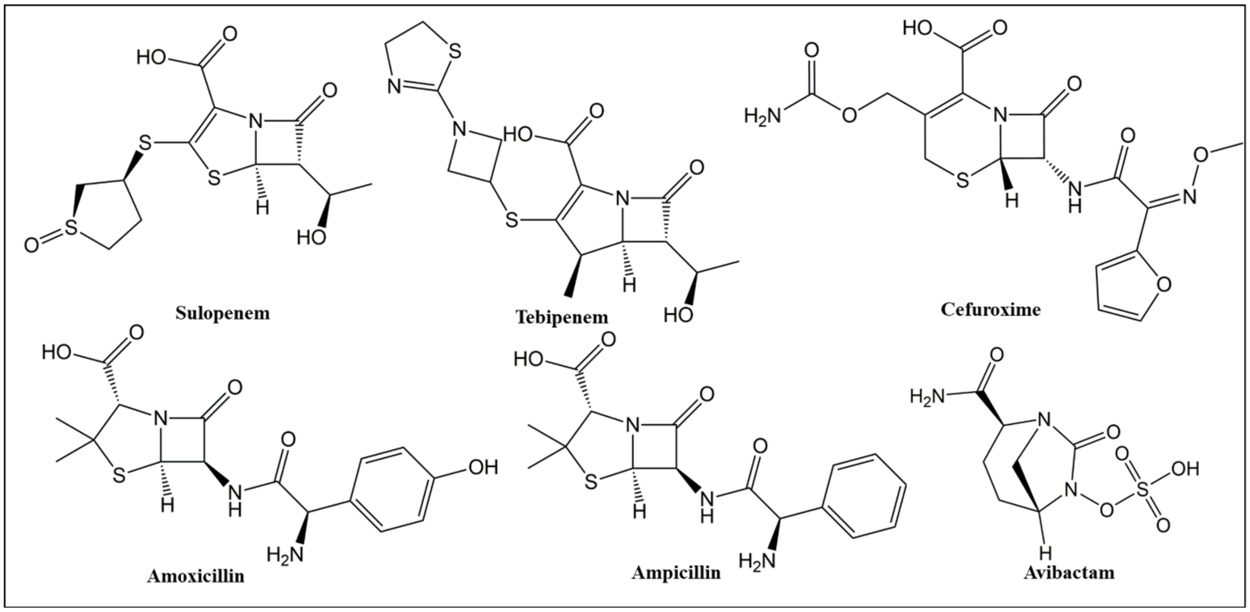


**Fig. S1.** Structures of β lactams SUP, TBP, CXM, AMX and AMP, and β-lactamase inhibitor AVI. Structures were derived from the PubChem database (https://pubchem.ncbi.nlm.nih.gov/) using the IUPAC name of the compounds.

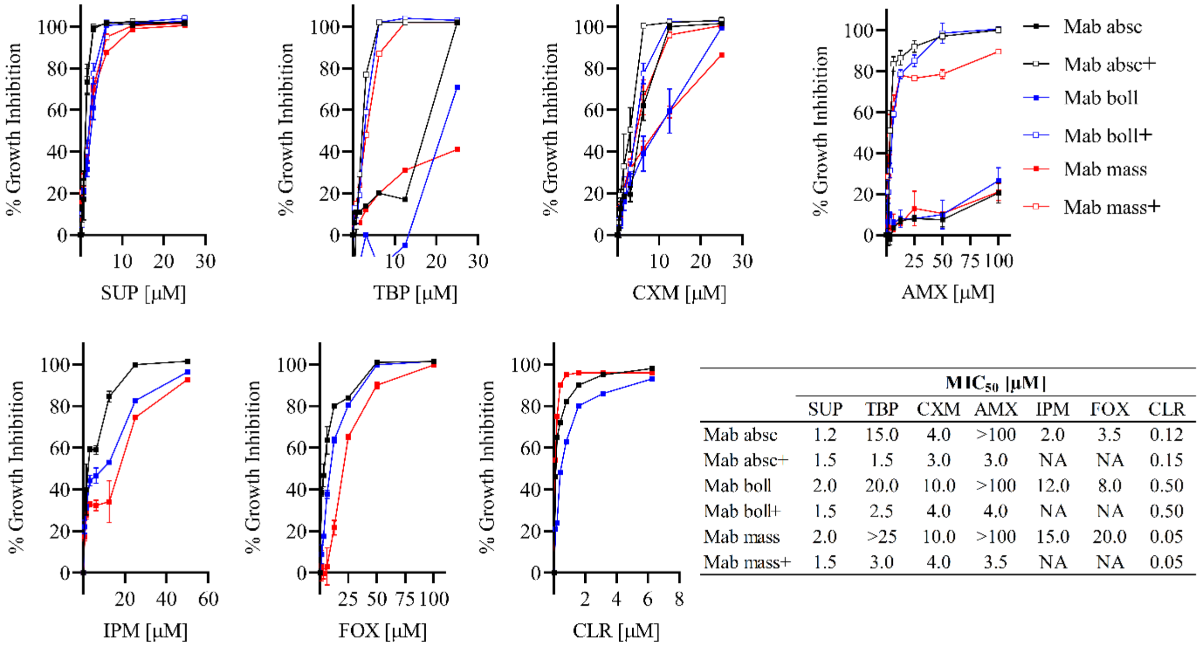


| | MIC$_{50}$ [μM] | | | | | | |
|---|---|---|---|---|---|---|---|
| | SUP | TBP | CXM | AMX | IPM | FOX | CLR |
| Mab absc | 1.2 | 15.0 | 4.0 | >100 | 2.0 | 3.5 | 0.12 |
| Mab absc+ | 1.5 | 1.5 | 3.0 | 3.0 | NA | NA | 0.15 |
| Mab boll | 2.0 | 20.0 | 10.0 | >100 | 12.0 | 8.0 | 0.50 |
| Mab boll+ | 1.5 | 2.5 | 4.0 | 4.0 | NA | NA | 0.50 |
| Mab mass | 2.0 | >25 | 10.0 | >100 | 15.0 | 20.0 | 0.05 |
| Mab mass+ | 1.5 | 3.0 | 4.0 | 3.5 | NA | NA | 0.05 |

**Fig. S2.** Growth inhibition dose-response curves for SUP, TBP, CXM and AMX with and without 4 ug/mL AVI against three *M. abscessus* subspecies reference strains. Mab absc, *M. abscessus* subsp. *abscessus* ATCC19977; Mab boll, *M. abscessus* subsp. *bolletii* CCUG50184T; Mab mass, *M. abscessus* subsp. *massiliense* CCUG48898T). '+', activity of β-lactam in the presence of 4 μg/mL AVI. CLR was included as assay control. IMP and FOX were included as clinically used parenteral comparators. The inserted table shows MIC$_{50}$ values (concentrations inhibiting 50% of growth) derived from the dose response curves. MIC values (concentrations inhibiting 90% of growth) derived from the curves are presented in Table 1. The experiments were carried out three time independently and means with standard deviations are shown.

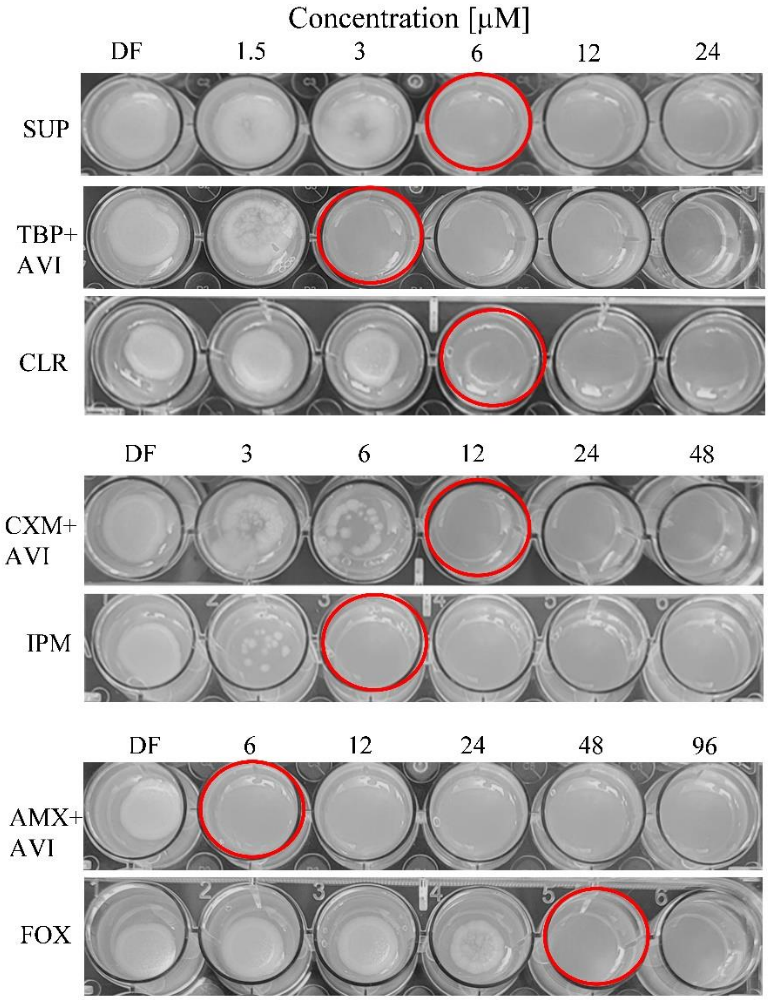


**Fig. S3.** Agar MIC of SUP, TBP+AVI, CXM+AVI and AMX+AVI for *M. abscessus* ATCC19977. $10^4$ CFU *M. abscessus* ATCC19977 culture were spotted on agar containing increasing β-lactam concentrations as indicated. '+AVI', 4 µg/mL AVI was included in the agar. The agar MIC (first concentration preventing visible growth), indicated by red circles, were SUP, 6 µM (2.5 µM); TBP+AVI, 3 µM (4.0 µM); CXM+AVI, 12 µM (5 µM); AMX+AVI, 6 µM (25 µM). Agar MIC for IPM and FOX, included as comparators, were 6 µM (20 µM) and 48 µM (30 µM). Agar MIC for CLR, included as assay control, was 6 µM (1.6 µM). Numbers in parentheses show MICs determined in liquid cultures (Table 1). The experiment was carried out twice, yielding similar results.
